# Dynamic partitioning shapes the *in vivo* organization of the *E. coli* RNA degradosome

**DOI:** 10.64898/2026.05.25.727690

**Authors:** Laura Troyer, Yu-Huan Wang, Kevin Wu, Seunghyeon Kim, Sangjin Kim

**Affiliations:** Department of Physics, University of Illinois Urbana-Champaign, Urbana, IL, USA; Center for Biophysics and Quantitative Biology, University of Illinois Urbana-Champaign, Urbana, IL, USA

## Abstract

The bacterial RNA degradosome is a central mediator of RNA turnover, yet how its components are organized and regulated in living cells remains unclear. Using live-cell single-molecule imaging, we quantified the spatial distribution and dynamics of the four major *Escherichia coli* degradosome components: RNase E, RhlB, PNPase, and enolase. These proteins occupied distinct membrane-associated and cytoplasmic pools whose relative abundance varied among proteins and changed with physiological state. In particular, PNPase underwent pronounced redistribution between membrane-associated and cytoplasmic states in response to altered RNA availability and growth conditions, whereas RNase E and RhlB remained largely membrane associated. To determine the functional consequences of this organization, we examined the degradation of *lacZ* reporter transcripts with different translation initiation strengths. PNPase and RhlB preferentially promoted degradation of weakly translated transcripts, whereas strongly translated transcripts were largely insensitive to their loss. Together, these results reveal that the bacterial RNA degradosome is not a static, uniformly assembled molecular machine. Instead, RNA decay is organized through dynamic intracellular partitioning of degradosome components, linking physiological state and translation status to transcript degradation.

## Introduction

RNA degradation is an important layer of bacterial gene regulation, controlling transcript abundance and contributing to stress responses and adaptation to environmental changes (1–3). As cellular conditions change, shifts in RNA synthesis and translation may impose different demands on RNA decay pathways. However, how bacterial RNA degradation machinery is organized and remodeled across physiological conditions remains unclear.

In bacteria, RNA decay is mediated by multienzyme degradosome assemblies, although their composition varies across species (4, 5). In *E. coli*, the RNA degradosome (RNAD) is organized around the essential endoribonuclease RNase E (RNE), which initiates degradation of most mRNAs and processes multiple classes of RNA (6, 7). RNE functions as a scaffold through interactions mediated by its carboxy-terminal domain (CTD) (5, 7). The CTD recruits the ATP-dependent DEAD-box family RNA helicase RhlB, the 3’→5’ exoribonuclease PNPase (PNP), and the glycolytic enzyme enolase (Eno) (6, 7). Together, these interactions are thought to organize RNA decay into a coordinated molecular machine.

RNAD is often depicted as a fully assembled molecular machine with fixed composition and stoichiometry (**Fig. 1**). Structural and biochemical studies have suggested that tetrameric RNE can simultaneously recruit multiple copies of RhlB, PNP, and Eno (4, 7, 8). This model has provided a useful framework for understanding degradosome architecture and function (7). However, whether this model accurately describes the organization of RNA decay machinery in living cells remains uncertain.

**Figure 1.**
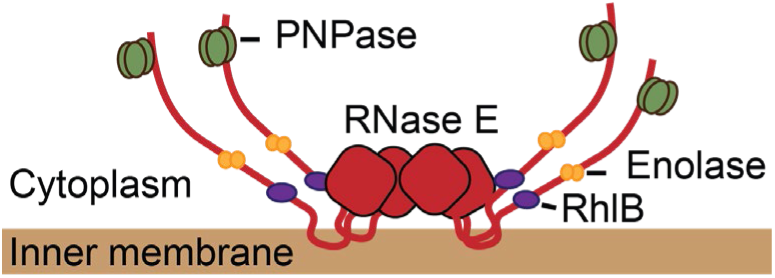
Schematic of the canonical *E. coli* RNA degradosome. Membrane-anchored RNase E (RNE) forms a tetrameric scaffold that recruits the accessory factors RhlB, PNP, and Eno to assemble the canonical membrane-associated RNA degradosome. In this model, RNase E is a tetramer (9), and each RNE monomer interacts with a monomeric RhlB (7, 10), a trimeric PNP (11), and a dimeric Eno (12).

Several observations suggest that RNAD composition may be more dynamic than the canonical model implies. Quantitative measurements indicate that RNAD components are not produced in stoichiometric ratios corresponding to a fully occupied scaffold (13, 14).

In addition, biochemical isolation of RNAD complexes has yielded widely varying estimates of component stoichiometry, with measured ratios depending strongly on purification conditions (10, 17, 18). These findings raise the possibility that substantial fractions of RNAD component proteins exist outside the canonical complex and that association with the RNE scaffold may be dynamically regulated rather than constitutive.

Recent advances in single-molecule imaging have begun to reveal the dynamic nature of bacterial RNA decay machineries. In *Bacillus subtilis*, proteins associated with the RNase Y-based degradosome exhibited multiple mobility states, and transcriptional inhibition altered both mobility and localization of those proteins (15). Such observations suggest that bacterial RNA decay machinery can undergo RNA-responsive reorganization. However, whether such reorganization reflects changes in scaffold composition, intracellular partitioning, or functional specialization of distinct component pools remains unclear.

In this study, we combined quantitative single-molecule imaging with functional measurements to define the *in vivo* organization of the *E. coli* RNAD. By resolving RNAD-associated and non-associated populations within intact cells, we determined how RNA availability and growth condition shape degradosome composition and intracellular partitioning. Using *lacZ* reporters with different ribosome binding strengths, we further assessed whether RhlB and PNP preferentially promote decay of weakly translated and prematurely terminated RNAs. Together, these approaches enabled us to move beyond the prevailing fully assembled complex model and identify dynamic intracellular partitioning as an organizing principle of bacterial RNA decay.

## Results

### Single-molecule imaging resolves distinct intracellular states of RNAD component proteins

To define the *in vivo* organization of the *E. coli* RNAD using single-molecule microscopy, we generated strains in which each of the four major RNAD components—RNE, RhlB, PNP, and Eno—was fused to the photoconvertible fluorescent protein mEos3.2 (19) and expressed from its native chromosomal locus (see **Table S1** for strain list and **Table S2** for strain construction). Using the single-particle tracking workflow previously established for RNE-mEos3.2 (20), we measured the localization and diffusion of individual molecules in living cells (**Fig. 2a**).

**Figure 2.**
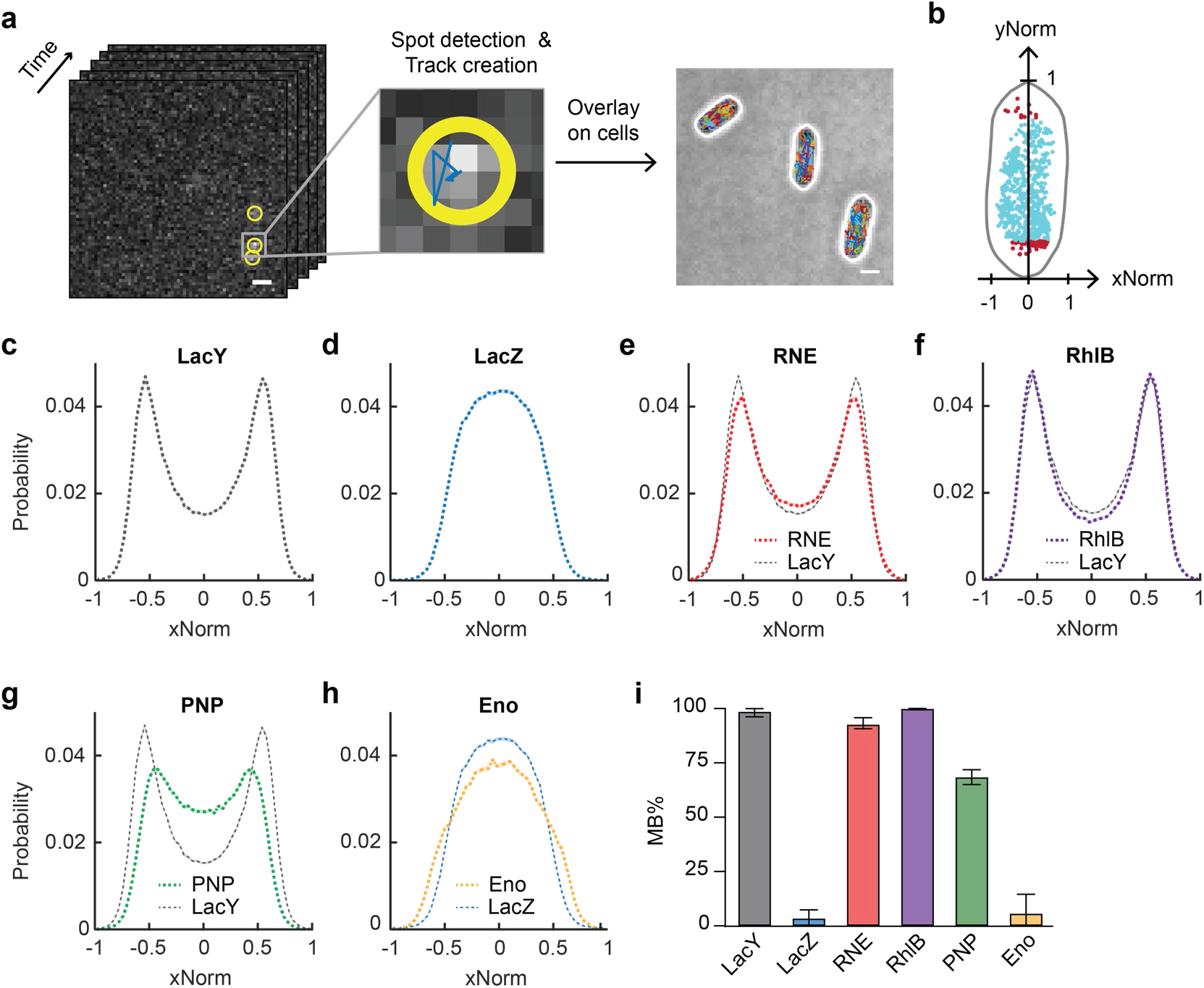
Subcellular localization of RNAD component proteins. (**a**) Microscopy and analysis workflow. Scale bar, 1 µm. (**b**) Example cell showing the localization of Eno-mEos3.2 in normalized cell axes. Cyan spots in the cylindrical region of the cell were included in the xNorm histogram, while red spots in the endcaps were excluded. (**c**–**h**) xNorm histograms for proteins imaged in live cells, including LacY (c), LacZ (d), RNE (e), RhlB (f), PNP (g), Eno (h). LacY and LacZ reference histograms are overlaid for comparison. SEM estimated by bootstrapping is shown as a shaded area. (**i**) Membrane-binding percentage (MB%) of the RNAD components and control proteins. Error bars represent 95% CIs estimated from the MCMC posterior distribution. Data statistics are provided in **Table S6**.

In our previous study, we showed that RNE is almost exclusively membrane bound (20). Because the RNAD is anchored to the inner membrane through the membrane-targeting sequence (MTS) of RNE (21), and because RhlB, PNP, and Eno lack intrinsic membrane-binding regions (14, 22, 23), membrane localization of these accessory factors provides a readout of their association with the RNE scaffold. We therefore inferred RNAD association from their partitioning between membrane and cytoplasmic pools, with membrane localization indicating the RNAD-associated population and cytoplasmic localization indicating a non-associated population. Furthermore, because a large membrane-associated degradosome complex is expected to diffuse slowly, we used diffusion as an independent and complementary measure of RNAD association.

### RNAD components partition differently between membrane-associated and cytoplasmic pools

Single-molecule localization was quantified using the position along the cell’s short axis, normalized by cell width (xNorm; **Fig. 2b**). In this analysis, membrane-localized molecules are enriched near the cell periphery, whereas cytoplasmic molecules show a more central distribution. As reference standards for these two localization patterns, we imaged LacY-mEos3.2 and LacZ-mEos3.2, which exhibited the expected membrane-like and cytoplasmic xNorm histograms, respectively (**Fig. 2c**,**d**). RNE and RhlB showed xNorm histograms very similar to that of LacY (**Fig. 2e**,**f**), indicating that both proteins were nearly fully membrane bound under these conditions. In contrast, the xNorm histogram of PNP differed from that of the membrane reference LacY, consistent with a substantial cytoplasmic population in addition to a membrane-associated population (**Fig. 2g**). Eno showed an xNorm histogram similar to that of LacZ (**Fig. 2h**), indicating that most Eno resided in the cytoplasm and was therefore not associated with RNAD. Thus, although all four proteins are considered canonical RNAD components, they occupy markedly different intracellular states in living cells.

We further quantified membrane association by fitting the xNorm histograms using the mathematical framework developed in our previous study (20). Model fitting was performed using a Markov chain Monte Carlo (MCMC) algorithm. The resulting membrane-bound fractions (MB%; **Fig. 2i**, **Fig. S1a**) were consistent with the qualitative xNorm patterns. In particular, PNP showed an MB% of 68% [65, 72], and Eno showed an MB% of 5.6% [0.2, 14]. Brackets indicate 95% credible intervals from the MCMC posterior distribution, hereafter referred to as 95% CIs.

### A common slow-diffusion state defines RNAD *in vivo*

We next analyzed the diffusion dynamics of RNAD components to determine whether they share a common mobility state corresponding to the membrane-associated degradosome. We calculated diffusion coefficients of individual proteins by fitting the mean-squared displacement (MSD) of each track to MSD(τ) = 4Dτ + b, where D is the diffusion coefficient, τ is the lag time (multiples of 21.7 ms), and b accounts for static and dynamic localization errors (24, 25). If multiple mobility populations are present, the resulting histogram of D is expected to reveal distinct slow- and fast-diffusion components.

The histograms of D_RNE_ and D_RhlB_ were well fit by a single Gaussian distribution with R^2^ > 0.97 (**Fig. 3a**,**b**), with the peak values of 0.014 µm^2^/s [0.013, 0.016] for RNE and 0.014 µm^2^/s [0.013, 0.015] for RhlB. We denote these fitted values as D_1,RNE_ and D_1,RhlB_, respectively. This result is consistent with the earlier MB% analysis, which suggested that most cellular RNE and RhlB are membrane bound, and therefore RNAD associated.

**Figure 3.**
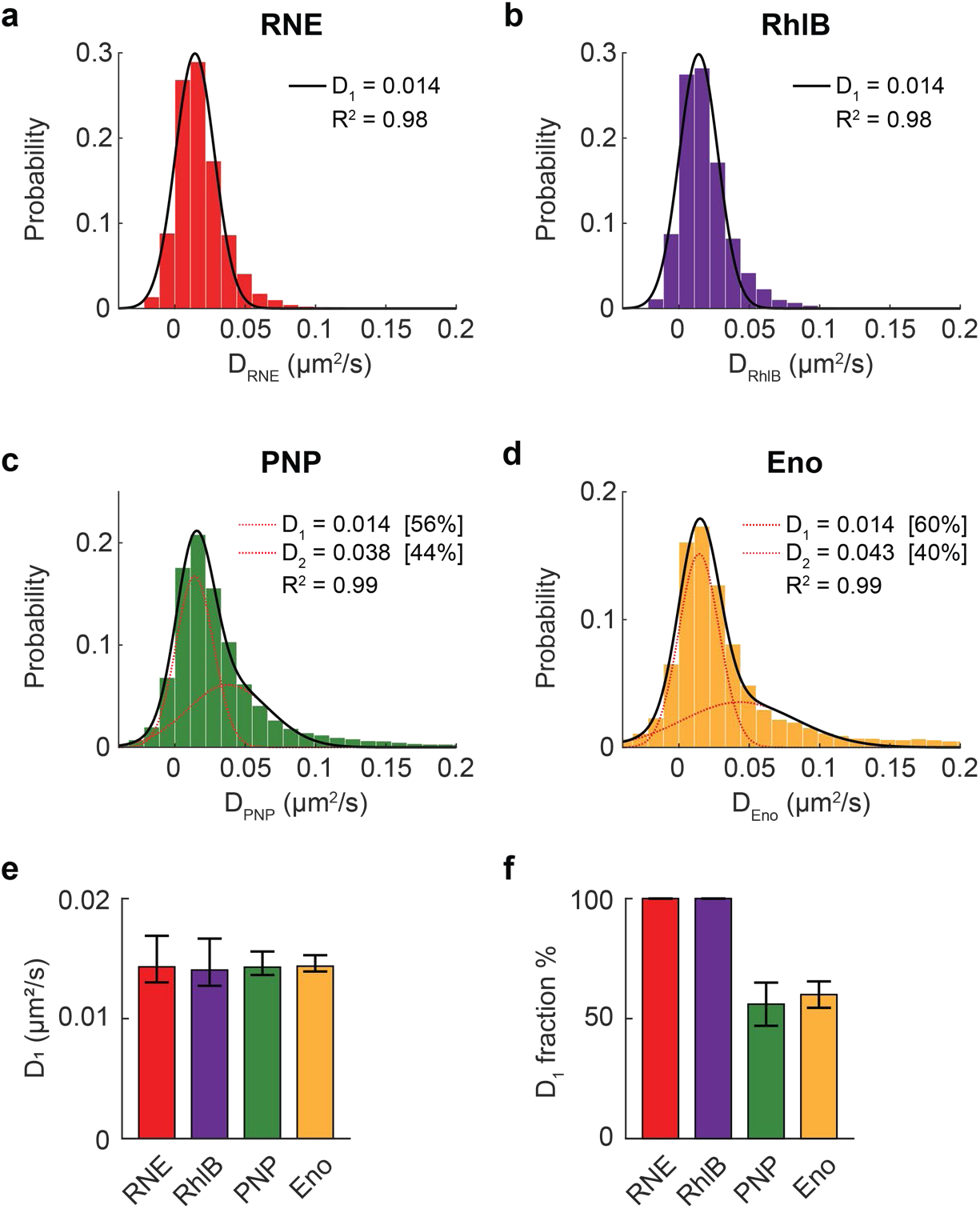
Mobility states of RNAD components. (**a-d**) Distributions of diffusion coefficients of RNE (a), RhlB (b), PNP (c), and Eno (d). Histograms were fit with either a single-Gaussian model, yielding D_1_ (a,b), or a two-Gaussian mixture model, yielding a slow and a fast diffusion coefficient D_1_ and D_2_, respectively (c,d). (**e**) D_1_ values obtained from the Gaussian fits in (a-d). Error bars indicate the 95% CI from fitting. (**f**) Fraction of molecules assigned to the slow-diffusion population from histogram fitting, with single-population fits treated as 100% slow. For two-population fits, error bars indicate 95% CIs from fitting. Detailed fitting results are provided in **Table S5**, and data statistics are provided in **Table S6**.

In contrast, the histograms of D_PNP_ and D_Eno_ contained pronounced long tails, indicating the presence of faster mobility populations (**Fig. 3c**,**d**). To separate these populations, the distributions were fit with a two-Gaussian model (R^2^ > 0.98), yielding a slow component (D_1_) and a fast component (D_2_). The slow components were close to those of RNE and RhlB with D_1,PNP_ = 0.014 µm^2^/s [0.014, 0.015] and D_1,Eno_ = 0.014 µm^2^/s [0.014, 0.015]. The fast components were D_2,PNP_ = 0.038 [0.032, 0.044] µm^2^/s and D_2,Eno_ = 0.043 [0.038, 0.049] µm^2^/s.

The common D_1_ value (∼0.014 µm^2^/s) across all four components defines a shared slow-diffusion state, consistent with the membrane-associated RNAD complex (**Fig. 3e**). The faster D_2_ populations of PNP and Eno likely represent cytoplasmic, non-RNAD-associated pools (**Fig. S2**). Approximately 50% of PNP and Eno molecules were assigned to this fast-diffusing state (**Fig. 3f**), indicating substantial partitioning away from the RNAD complex.

Notably, the slow D_1_ fraction of Eno (∼50%) was higher than the MB% estimated from the xNorm histogram (∼6%), contrary to the expectation that the slow fraction would largely correspond to the membrane-bound population. This discrepancy suggests that the very fast-moving fraction of Eno may be underrepresented in the diffusion analysis. Although both analyses used the same live-cell imaging dataset, diffusion analysis required trajectories of at least 12 frames for robust D estimation, likely biasing the dataset toward slower (and longer) trajectories. In contrast, the xNorm analysis included all trajectories, including those shorter than 12 frames, and was therefore able to capture rapidly diffusing cytoplasmic molecules. Consistent with this interpretation, restricting the xNorm analysis to the trajectories used for diffusion analysis increased the apparent MB% of Eno (**Fig. S3**). Thus, for Eno, the exact value of MB% depends in part on trajectory-selection criteria. For subsequent analyses, we used MB% values estimated from the full dataset to ensure inclusion of rapidly diffusing molecules.

### Membrane-associated pools reveal near-canonical RNAD stoichiometry under the reference condition

The differential partitioning of RNAD components raised the question of how closely the membrane-associated population resembles the canonical degradosome complex *in vivo*. To estimate RNAD stoichiometry under the reference condition, we combined the membrane-bound fractions measured by live-cell xNorm analysis with published protein abundance values for *E. coli* derived from ribosome profiling (13), while noting that these abundance measurements were obtained under different growth conditions.

Because ribosome profiling reports synthesis of individual polypeptide chains, the published abundance values correspond to monomeric subunits. We therefore first converted these values to functional oligomeric forms for tetrameric RNE (RNE_4_), trimeric PNP (PNP_3_), and dimeric Eno (Eno_2_). We then multiplied them by the corresponding MB% estimated from xNorm histograms (**Fig. 2i**). This yielded an ensemble-average membrane-associated ratio of approximately 1 RNE_4_ : 4-5 RhlB : 3-4 PNP_3_ : 2 Eno_2_ (**Table 1**). Thus, RhlB and PNP were present in the membrane-associated pool at levels broadly consistent with the canonical expectation for a fully assembled RNAD complex, 1 RNE_4_: 4 RhlB: 4 PNP_3_: 4 Eno_2_ (**Fig. 1**), while Eno appeared lower.

**Table 1.** Estimated membrane (MB)-associated stoichiometry of RNAD components. Published protein abundance values derived from ribosome profiling (13) were converted to functional oligomeric forms and multiplied by the corresponding membrane-bound fraction (MB%) measured from xNorm histogram fitting. MB% values are shown as the mean ± s.d. of the MCMC posterior distribution. MB-associated abundances were normalized to RNE, and errors represent uncertainty propagated from MB% estimation.

|  | MB% | Abundance from EZRDM |  | Abundance from MOPS glucose |  |
| --- | --- | --- | --- | --- | --- |
|  |  | Multimer | MB-associated multimers, relative to RNE | Multimer | MB-associated multimers, relative to RNE |
| RNE<sub>4</sub> | 92.6 ± 1.2 | 682 | 1 | 265 | 1 |
| RhIB | 99.8 ± 0.2 | 3,459 | 5.5 ± 0.1 | 1,055 | 4.3 ± 0.1 |
| PNP<sub>3</sub> | 68.3 ± 1.9 | 3,968 | 4.3 ± 0.1 | 1,181 | 3.3 ± 0.1 |
| Eno<sub>2</sub> | 5.6 ± 3.8 | 27,111 | 2.4 ± 1.6 | 10,617 | 2.4 ± 1.7 |

One caveat is that our MB% values were measured in cells grown in M9 minimal medium supplemented with 0.2% glycerol, casamino acids, and thiamine (M9 gly + CAAT) at 30°C, whereas the protein abundance values were obtained from cells grown in Neidhardt EZ rich defined medium or MOPS medium with 2% glucose at 37°C. Despite this difference in growth conditions, the inferred membrane-associated stoichiometry places RhlB and PNP broadly near the canonical expectation. Eno appeared lower than the canonical value of 4, although this estimate is more sensitive to uncertainty in MB% because of the large cellular abundance of Eno. Indeed, the uncertainty range arising from MB% estimation spans the canonical value.

Together, this analysis reconciles the canonical degradosome model with the single-cell partitioning data by showing that a near-canonical membrane-associated complex can coexist with large non-associated pools of specific accessory factors.

### RNA depletion selectively remodels PNPase and enolase localization

We next asked whether this partitioning is regulated by RNA availability. To reduce RNA levels, cells were treated with rifampicin (rif), an inhibitor of transcription initiation (26), for 15 min before imaging, and were imaged within 1 hour in the continued presence of rif on the agarose pad. We also imaged LacY and LacZ under the same conditions as reference proteins for 100% and 0% membrane binding, respectively, to account for any rif-induced changes in xNorm distributions (**Fig. 4a**,**b**).

**Figure 4.**
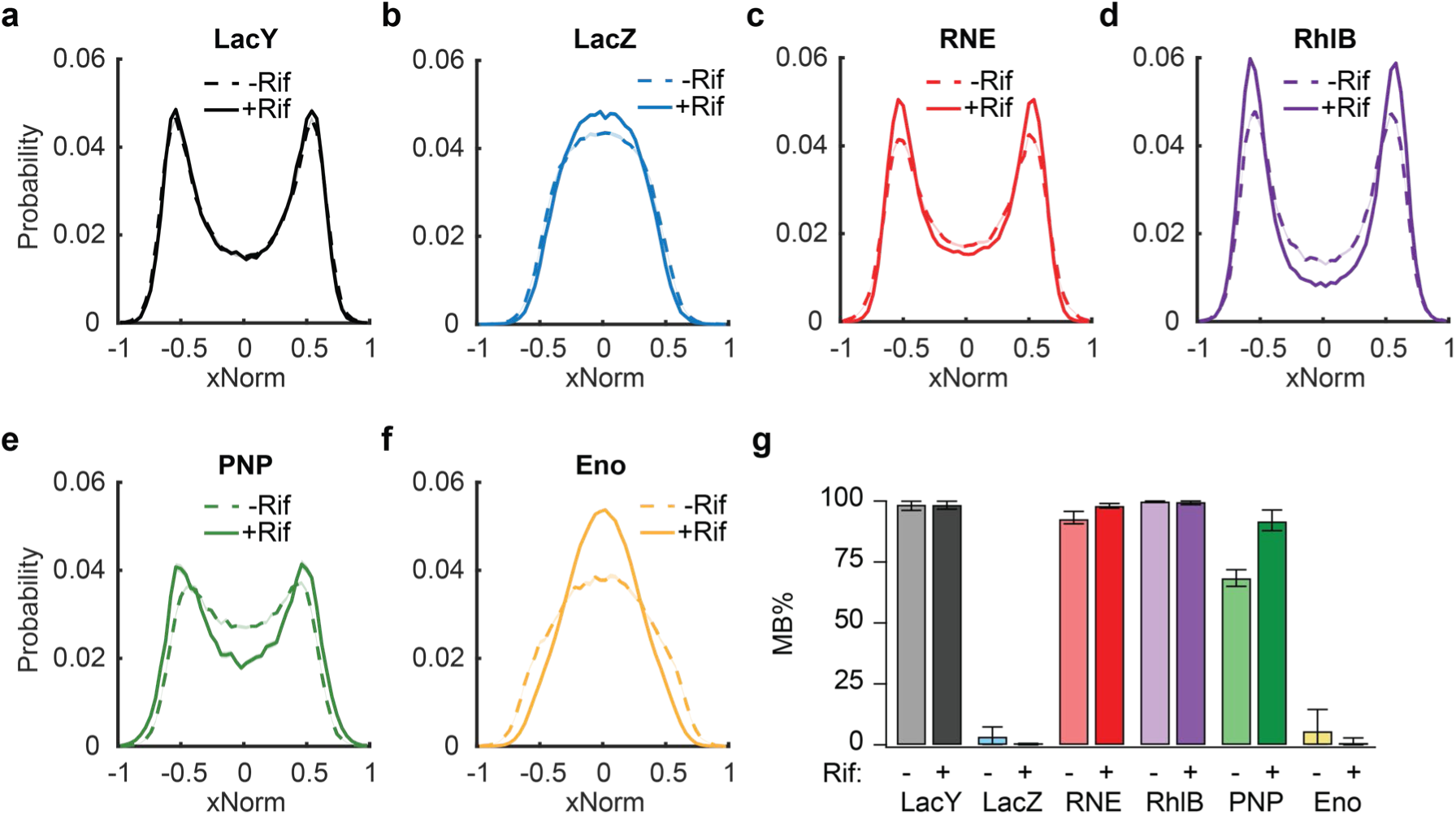
RNAD component localization in rifampicin-treated cells. (**a**-**b**) Reference proteins: LacY, a membrane protein, and LacZ, a cytoplasmic protein. (**c**-**f**) The four canonical RNAD components. Shaded colored regions around the xNorm histogram lines represent the SEM estimated by bootstrapping. (**e**) Membrane-binding percentage (MB%) of the RNAD components and control proteins before and after rifampicin treatment. Error bars represent 95% CIs estimated from the MCMC posterior distribution. Detailed data statistics are reported in **Table S6**.

Surprisingly, the RNAD components exhibited different localization responses upon rif treatment (**Fig. 4c**-**f**, **Fig. S1b**, **Fig. S4a**). Notably, RNE and RhlB were largely unaffected by rif treatment and remained predominantly membrane localized, whereas PNP became more membrane-associated and Eno became more cytoplasmic. Quantification of MB% by xNorm fitting confirmed these trends. PNP displayed a marked increase in MB%, from 68% to 92%, indicating a reduction in its cytoplasmic pool (**Fig. 4e**). The rif-induced increase in membrane association of PNP was also evident in two-dimensional localization histograms (**Fig. S5**), in which a central depletion region became more apparent with higher MB%.

This redistribution also altered the inferred composition of the membrane-associated RNAD pool. Assuming that total protein abundance does not change substantially during the short rif-treatment period, we applied the MB% values measured after rif treatment to the same abundance-based framework used above. This analysis indicated that the inferred PNP occupancy increased, reaching approximately 5.4 PNP_3_ per RNE_4_ when combined with abundance values from EZRDM and 4.2 PNP_3_ per RNE_4_ when combined with abundance values from MOPS glucose (**Table 2**). In contrast, Eno was strongly depleted from the membrane-associated pool, with an inferred value of approximately 0.3 Eno_2_ per RNE_4_ in both abundance datasets.

**Table 2.** Estimated membrane (MB)-associated stoichiometry of RNAD components in rifampicin-treated cells. As in Table 1, published protein abundance values from ribosome profiling (13) were converted to functional oligomeric forms and multiplied by the membrane-bound fraction (MB%) measured from rif-treated cells. MB% values are shown as the mean ± s.d. of the MCMC posterior distribution. MB-associated abundances were normalized to RNE, and errors represent uncertainty propagated from MB% estimation.

|  | MB% | Abundance from EZRDM |  | Abundance from MOPS glucose |  |
| --- | --- | --- | --- | --- | --- |
|  |  | Multimer | MB-associated multimers, relative to RNE | Multimer | MB-associated multimers, relative to RNE |
| RNE<sub>4</sub> | 98.0 ± 0.5 | 682 | 1 | 265 | 1 |
| RhIB | 99.6 ± 0.4 | 3,459 | 5.2 ± 0.0 | 1,055 | 4.0 ± 0.0 |
| PNP<sub>3</sub> | 91.7 ± 2.7 | 3,968 | 5.4 ± 0.1 | 1,181 | 4.2 ± 0.1 |
| Eno<sub>2</sub> | 0.7 ± 0.7 | 27,111 | 0.3 ± 0.3 | 10,617 | 0.3 ± 0.3 |

Together, these findings reveal an RNA-dependent spatial reorganization of specific RNAD components. In particular, PNP is slightly underrepresented in the membrane-associated RNAD pool under untreated conditions, whereas rif treatment shifts PNP closer to canonical occupancy. One possible explanation is that abundant cytoplasmic RNA promotes retention of PNP in the cytoplasm, whereas RNA depletion after rif treatment shifts the equilibrium toward association with the membrane-bound RNE scaffold.

### RNAD organization shifts in succinate-grown cells

Cellular RNA abundance and transcriptional flux vary substantially across growth conditions (27, 28). We therefore asked whether RNAD organization can be modulated under distinct physiological states. To test this, we analyzed diffusion and localization of RNAD components in cells grown in M9 succinate medium without CAAT. Succinate is a poorer carbon source than glycerol under our conditions, with doubling times of 153 min for MG1655 grown in M9 succinate and 89 min in M9 gly + CAAT at 30°C.

Single-molecule tracking revealed diffusion behavior qualitatively similar to that observed in M9 gly + CAAT. The histograms of D_RNE_ and D_RhlB_ were well fit by a single population, whereas those of PNP and Eno exhibited two mobility populations (**Fig. S6**). All components shared a common slow-diffusion state with D_1_ of ∼0.02 µm^2^/s, consistent with RNAD association (**Fig. S6**). These D_1_ values were 1.5-2-fold higher than those measured in M9 gly + CAAT, indicating increased mobility of the RNAD-associated state in cells grown in M9 succinate. The fast-diffusing populations of PNP and Eno (D_2_) were also shifted to higher values relative to M9 gly + CAAT (**Fig. S6**).

We next examined xNorm histograms. Most proteins, including LacY and LacZ, showed little change relative to M9 gly + CAAT. PNP shifted toward a slightly higher MB%, whereas Eno shifted toward a lower MB% (**Fig. 5**, **Fig. S1c**, **Fig. S4b**), a trend qualitatively similar to that observed after rif treatment (**Fig. 4e**,**f**).

**Figure 5.**
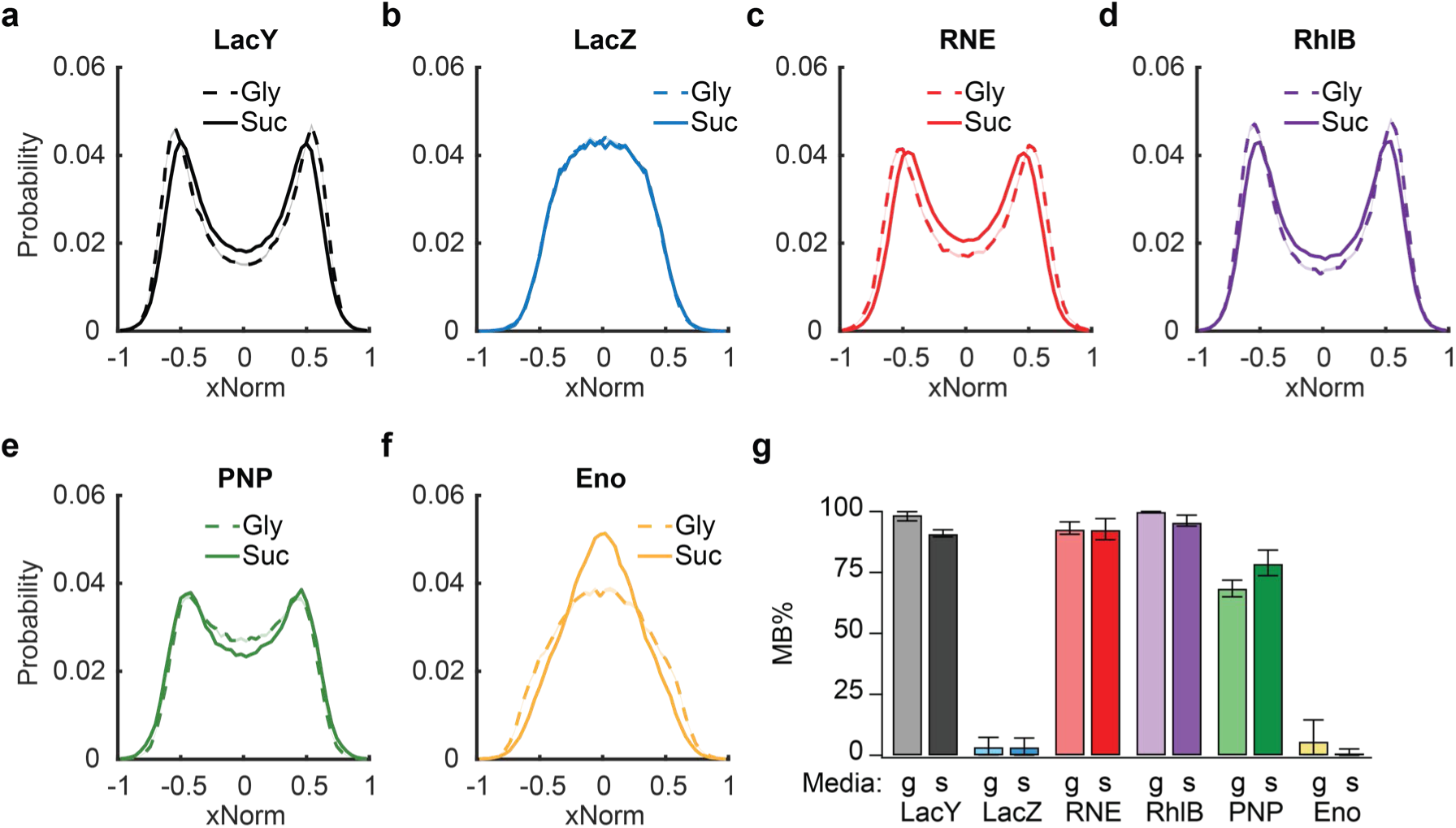
Subcellular localization of RNAD components in cells grown in M9 succinate. (**a-f**) xNorm histograms of proteins in cells grown in M9 succinate, including reference proteins LacY (a) and LacZ (b) as well as the four RNAD component proteins, RNE (c), RhlB (d), PNP (e), and Eno (f). Shaded colored regions around the xNorm histogram lines represent the SEM from bootstrapping. Data from M9 glycerol + CAAT are drawn for a comparison. (**g**) MB% quantified from the xNorm histograms shown in (a-f). ‘g’ represents M9 glycerol +CAAT and ‘s’ represents M9 succinate. Error bars represent 95% CIs estimated from the MCMC posterior distribution. Data statistics are listed in **Table S6**.

These observations indicate that RNAD organization is not constant across growth conditions. Instead, both the localization and mobility of degradosome components are remodeled by physiological state. One possible interpretation is that these changes reflect altered partitioning of RNAD components in response to differences in RNA availability and carbon metabolism across growth conditions. Because glycerol and succinate enter different points in central carbon metabolism (29), enolase partitioning may be linked to changes in metabolic state, potentially including altered metabolic flux and increased coupling between glycolytic and RNA decay pathways under carbon-limited conditions (30). Also, these observations raise the possibility that enolase function may depend in part on its partitioning between cytoplasmic and membrane-associated RNAD pools.

### RhlB and PNP preferentially promote decay of weakly translated *lacZ* transcripts

The RNA-dependent redistribution of PNP led us to a functional question: Do RNAD components contribute differently to decay of distinct classes of RNA substrates? In our recent work, we showed that translation initiation rate, altered by ribosome binding site (RBS) strength, influences the abundance of prematurely released transcripts arising from disrupted transcription-translation coupling (31). Based on this, we asked whether RhlB and PNP contribute more strongly to the decay of weakly translated mRNAs than of efficiently translated transcripts.

To test this idea, we examined the degradation kinetics of *lacZ* mRNA carrying either a strong (native) or weak RBS (**Fig. 6a**). In our previous work, 75-s transient induction of *lacZ* and time-course quantification of *lacZ* 5’ levels by reverse transcription quantitative polymerase chain reaction (RT-qPCR) resolved two decay phases: *k*_d1_, reflecting decay of nascent or prematurely released transcripts, and *k*_d2_, corresponding to decay of fully transcribed transcripts (31). *lacZ* transcripts bearing the native RBS exhibited negligible *k*_d1_, indicating minimal co-transcriptional decay, minimal premature release, and predominantly post-transcriptional decay (**Fig. 6b**). In contrast, *lacZ* transcripts with a weak RBS displayed an elevated *k*_d1_, consistent with increased production of prematurely released RNA species before the *lacZ* 3’ signal increased (**Fig. 6c**).

**Figure 6.**
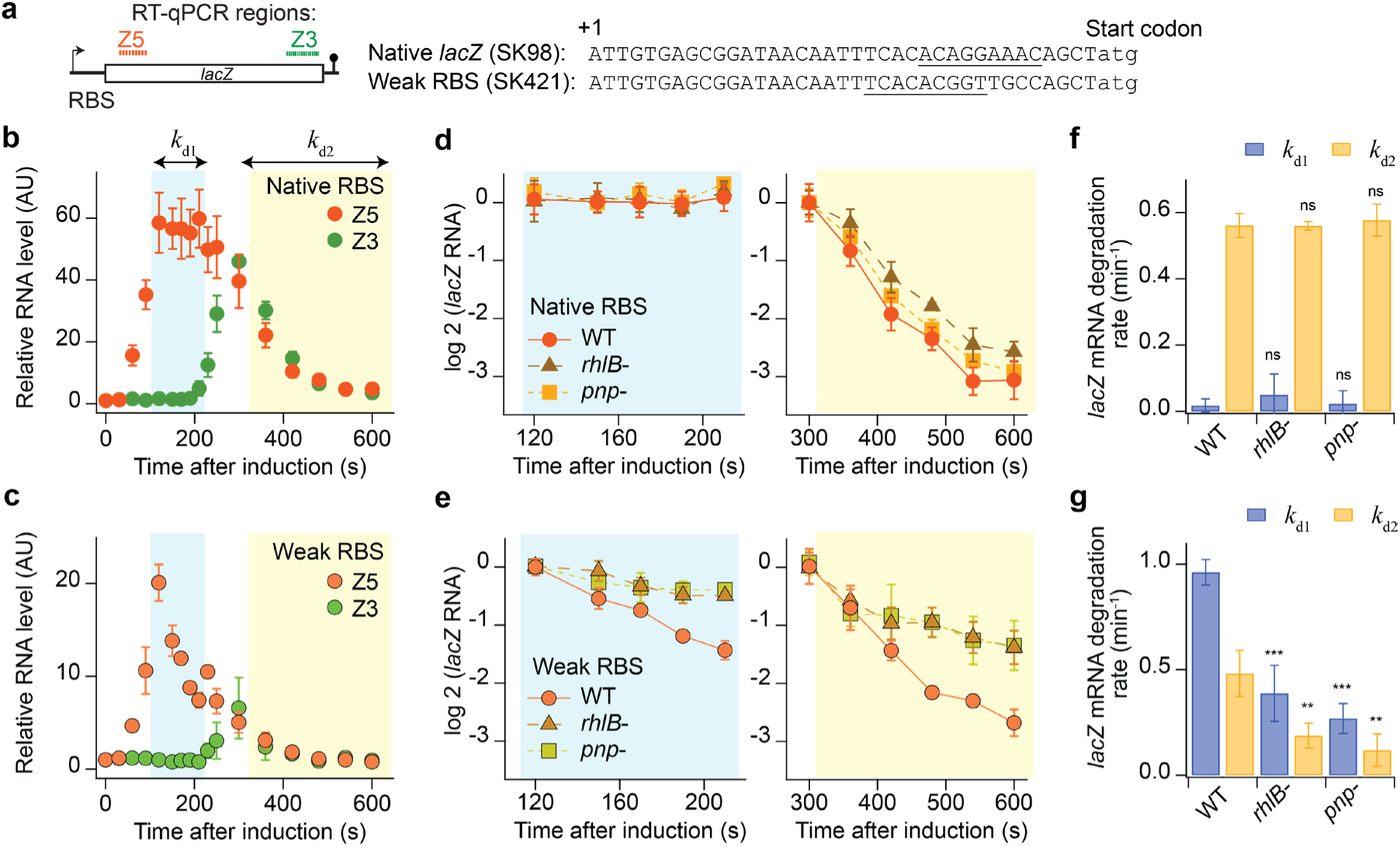
mRNA degradation analysis using the *lacZ* reporter in wild-type, *rhlB*-, and *pnp*- strains. (**a**) Schematic of the *lacZ* reporter at its native chromosomal locus with different ribosome-binding-site (RBS) sequences. mRNA sequences are shown from the first base (+1) to the start codon (atg). Underlined sequences are Shine-Dalgarno elements estimated by an RBS calculator (32). The 5’ and 3’ RT-qPCR amplicons are indicated as Z5 and Z3, respectively. (**b**,**c**) Relative RNA levels measured after induction with 0.2 mM IPTG at t = 0 and re-repression with 500 mM glucose at t = 75 s for reporters containing the native RBS (SK98) or a weak RBS (SK421). Blue and yellow shaded regions indicate the time windows used to quantify the Z5 decay rates, *k*_d1_ and *k*_d2_, respectively, by exponential fitting. (**d**,**e**) Changes in *lacZ* Z5 RNA levels within the blue and yellow time windows, respectively, for the native-RBS reporter (d) and the weak-RBS reporter (e), normalized to the first time point shown in each graph. (**f**,**g**) Decay rates *k*_d1_ and *k*_d2_ obtained from the data in (d) and (e), respectively. Native RBS strains are SK98 (WT), SK301 (*rhlB*-), and SK306 (*pnp*-) (f), and weak RBS strains are SK421 (WT), SK437 (*rhlB*-), and SK431 (*pnp*-) (g). Data are mean ± s.d. of four (SK98, SK421, SK301, SK437) and three (SK306, SK431) biological replicates. In panel f, *k*_d1_, *P* = 0.36 (SK301), *P* = 0.83 (SK306); *k*_d2_, *P* = 0.95 (SK301), *P* = 0.64 (SK306), from two-sided two-sample *t*-tests relative to SK98. In panel g, *k*_d1_, \*\*\**P* = 0.0010 (SK437), \*\*\**P* = 0.00020 (SK431); *k*_d2_, \*\**P* = 0.0056 (SK437), \*\**P* = 0.0092 (SK431), from two-sided two-sample *t*-tests relative to SK421.

To determine whether RhlB and PNP contribute functionally to mRNA decay, we measured *lacZ* mRNA degradation kinetics in wild-type, Δ*rhlB*, and Δ*pnp* strains. For *lacZ* transcripts with the native RBS, *k*_d1_ remained close to zero, and the major decay phase (*k*_d2_) was indistinguishable between the wild-type and deletion strains, consistent with RNE-mediated cleavage remaining rate-limiting for this transcript (**Fig. 6d**,**e**).

In contrast, for *lacZ* transcripts containing a weak RBS, relative RNA levels declined more slowly in Δ*rhlB* and Δ*pnp* strains than in the wild-type strain in the *k*_d1_ and *k*_d2_ fitting regions (**Fig. 6f**), resulting in reductions in both *k*_d1_ and *k*_d2_ (**Fig. 6g**). *k*_d1_ was particularly sensitive to loss of RhlB and PNP (**Fig. 6g**). These results indicate that RhlB and PNP contribute to efficient degradation of weakly translated or non-translating transcripts, whereas decay of strongly translated transcripts is less dependent on these accessory components.

## Discussion

By combining live-cell single-molecule localization, diffusion measurements, and functional RNA decay assays, we examined how the major components of the *E. coli* RNA degradosome (RNAD) are organized *in vivo*. Our results support a view in which RNAD is a membrane-anchored but dynamically partitioned system composed of component-specific membrane-associated and cytoplasmic pools whose balance shifts with physiological state.

This view helps bridge a long-standing gap between biochemical models of the RNAD and its behavior in living cells. Earlier studies based on affinity purification yielded differing pictures of accessory-protein occupancy, with some suggesting over-representation of RhlB (10, 17) and Eno (17) and under-representation of PNP (10, 17, 18), relative to canonical stoichiometry. Our approach does not resolve the composition of individual complexes directly, but it does provide an *in situ* readout of how RNAD components partition between membrane-associated and cytoplasmic pools. This distinction reconciles the canonical degradosome model with the single-molecule localization and partitioning data by showing that a near-canonical membrane-associated complex can coexist with large non-associated pools of specific accessory factors.

Among the accessory factors, PNP showed the clearest evidence of dynamic partitioning. In untreated cells, a substantial fraction of PNP resided in the cytoplasm, but this fraction decreased after rif treatment and in succinate-grown cells. One plausible interpretation is that abundant cytoplasmic RNA favors RNA-bound pools of PNP, whereas RNA depletion shifts the equilibrium toward PNP association with the membrane-bound RNE scaffold. In this view, PNP may interact with RNA in both scaffold-associated and non-associated states, with the balance between these pools set by cellular RNA availability. This dynamic behavior may also explain why biochemical studies have not converged on a single degradosome stoichiometry in the literature. If a substantial fraction of PNP is conditionally associated with RNE, its apparent occupancy will depend not only on growth condition but also on how complexes are extracted or stabilized.

A possible functional consequence of this organization is a division of labor between membrane-associated decay initiation and downstream processing of RNA fragments. RNE is anchored to the inner membrane and initiates RNA decay by endonucleolytic cleavage. PNP, in contrast, partitions between the membrane-associated scaffold and the cytoplasm, allowing it to act on RNA decay intermediates either locally at the RNAD or after their release from the scaffold (**Fig. 7**). Maintaining a substantial cytoplasmic pool of PNP could therefore facilitate clearance of decay intermediates without requiring all processing to occur at the membrane-bound scaffold. At the same time, scaffold association of PNP likely remains important for coordinated decay of specific substrates, enabling rapid local handoff after RNE-mediated cleavage (33). This view is consistent with recent 3’-end profiling showing that most *E. coli* mRNAs are decay intermediates rather than full-length transcripts, that these intermediates are depleted from ribosome-rich fractions, and that they accumulate further in Δ*pnp* cells (34). It is also consistent with transcriptome-wide studies showing that different classes of *E. coli* mRNAs respond differently to mutations in RNAD components, suggesting that RNA decay need not proceed through a single uniformly assembled degradosome pathway (35).

**Figure 7.**
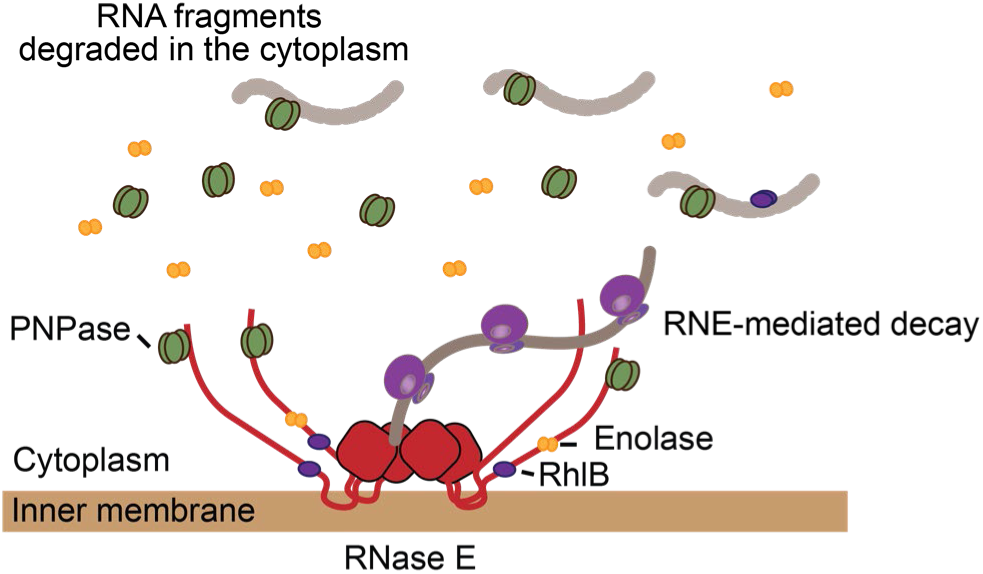
Model for dynamic partitioning and functional organization of RNAD components in *E. coli*. RNE is anchored to the inner membrane and serves as a scaffold for assembly of the RNAD. RhlB is largely associated with the membrane-bound scaffold, whereas PNP partitions between membrane-associated and cytoplasmic pools in an RNA-dependent manner. Eno is largely cytoplasmic, with only a smaller fraction associated with the membrane-bound complex. In this model, membrane-associated RNE initiates RNA decay by endonucleolytic cleavage, after which RNA fragments can be further degraded by PNP either at the scaffold or in the cytoplasm. Grey lines represent RNA.

The *lacZ* decay experiments add an important functional dimension to this model. Loss of *rhlB* or *pnp* had little effect on degradation of strongly translated *lacZ* transcripts, but both deletions slowed decay of weakly translated transcripts. In particular, *k*_d1_, reflecting decay of prematurely terminated transcripts (31), showed notable sensitivity. Thus, RhlB and PNP matter most when translation is weak and transcripts are likely to be prematurely released or less covered by ribosomes. Although these experiments do not directly assign different RNA substrates to cytoplasmic and membrane-associated pools, they support the broader idea that degradosome factors contribute unequally to RNA decay across substrate classes, providing a functional counterpart to their differential intracellular partitioning.

Several caveats are worth keeping in mind. Our stoichiometry estimates combine measured MB% values with published abundance data rather than direct copy-number measurements under the same growth conditions. In addition, the relevant oligomeric state of RhlB *in vivo* remains uncertain. We assumed monomeric RhlB interacting with one RNE monomer based on ref (7, 10); however, other studies have suggested that RhlB may function as a dimer (36, 37). If a dimeric RhlB interacts with each RNE monomer, the inferred RhlB occupancy would be reduced (i.e. under occupied). These limitations affect the precision of the stoichiometric estimates, but not the central conclusions from the imaging data: RhlB is strongly membrane associated, PNP is dynamically partitioned, and Eno is only weakly represented in the membrane-associated pool.

Overall, our results support a view of the *E. coli* RNA degradosome as a membrane-anchored but compositionally flexible RNA decay system. Rather than existing as one static assembly, it appears to comprise component-specific associated and non-associated populations whose balance shifts with RNA availability and growth condition. More broadly, our findings suggest that bacterial cells can regulate RNA decay not only through enzyme activity or substrate recognition, but also by redistributing decay factors between distinct intracellular pools. Given that RNE serves as a scaffold for RNA degradosome assembly across bacteria and chloroplasts (5), dynamic intracellular partitioning may represent a broader principle for organizing RNA decay machinery *in vivo*.

## Materials and Methods

### Cloning and strain construction

Strains used in this study are listed in **Table S1** with the details of construction listed in **Table S2**. Doubling times were measured as described previously(20). Briefly, cells were grown overnight in M9 glycerol +CAAT or in M9 succinate at 30°C, diluted 10^4^-fold, and the optical density at 600 nm (OD_600_) was measured using a microplate reader (Synergy HTX multi-mode reader, BioTek), with continuous shaking. Data analysis included fitting the background-subtracted data to an exponential function within OD_600_ between 0.001 to 0.1 (**Table S3**).

### Cell growth and sample preparation

Cells were grown in M9 minimal media supplemented with 0.2% (v/v) glycerol (Invitrogen 15514011), 0.1% (w/v) casamino acids (Bacto 223050), and 1 µg/mL thiamine (Research Products International T21020) at 30°C with shaking at 220 rpm. Starter cultures were inoculated from several colonies and grown for 6–8 h, then diluted ≥3000-fold and grown overnight to early exponential phase (OD_600_ = 0.15–0.25) prior to imaging.

For M9 succinate conditions, cells were grown in M9 minimal media supplemented with 0.34% (w/v) sodium succinate (Sigma-Aldrich S2378) at 30°C with shaking at 220 rpm. Starter cultures were grown for 12–72 h, diluted ≥1000-fold, and then grown for several days to early exponential phase (OD_600_ = 0.095–0.105) prior to imaging.

Glass slides (Fisher 12-544-1) and #1.5 glass coverslips (Fisher 12544A or VWR 16004-344) were cleaned by three rounds of sonication (15 min each) in 100% ethanol, ∼80% ethanol, and Milli-Q water. Cleaned slides and coverslips were stored in Milli-Q water until the day of imaging and dried with nitrogen gas immediately before use.

Agarose pads (1% w/v) were prepared by dissolving agarose (Invitrogen 16500-100) in the same type of medium used for cell growth. Molten agarose was solidified between two cleaned glass slides to ensure a flat surface. Cells were applied to the agarose pad, and excess liquid was removed by air drying for ∼1 min. A cleaned coverslip was then placed on top and sealed with VALAP. Samples were imaged immediately. When rifampicin-treated cells were imaged, 200 ug/mL of rifampicin was added to the molten agarose kept at 60°C and solidified right away.

### Drug treatment

Cells were treated with rifampicin (400 μg/mL, Sigma R3501) for 15 min in the liquid culture with shaking at the growth temperature.

### Single-molecule imaging

Bright-field and single-molecule mEos3.2 fluorescence imaging was performed on a Nikon Ti-2 microscope as previously described (20). Imaging conditions, including hardware configuration and laser settings, were identical to those used in our previous study (20). Briefly, a quasi-TIRF illumination geometry was used, with a 405 nm laser for photoconversion of mEos3.2 and a 561 nm laser for excitation of the photoconverted state. Images were acquired at a frame interval of 21.7 ms under continuous illumination for 3 min. Each strain was imaged in three biological replicates, with 6-16 movies collected per replicate.

### *lacZ* induction, RNA extraction, and RT-qPCR

Pulsed induction of the *lacZ* reporter, RNA extraction, and RT-qPCR were performed as previously described (31). Cells were grown in M9 minimal medium supplemented with 0.2% (v/v) glycerol and CAAT at 30°C to early exponential phase (OD_600_ = 0.2). Following induction with 0.2 mM IPTG at t = 0, 0.4 mL of culture was withdrawn every 20-60 s and immediately mixed with an equal volume of pre-cooled RNAlater Stabilization Solution (Life Technologies). 500 mM glucose was added to the culture at t = 75 s to repress the transcription. After mixing with RNAlater, samples were incubated on ice for 20 min and then centrifuged. Cell pellets were resuspended in lysozyme solution (10 mg/mL of lysozyme in 10 mM Tris-HCl (pH 8) with 1 mM EDTA). Total RNA was extracted using the PureLink RNA Mini Kit (Life Technologies) with on-column DNase treatment according to the manufacturer’s instructions. RT-qPCR was performed using KAPA SYBR FAST qPCR Master Mix (KAPA Biosystems) on a Bio-Rad CFX Connect system. Primer sequences are provided in **Table S4**.

Relative RNA levels were calculated as fold change relative to baseline (pre-induction samples) using the 2^-ΔCt^ method, where Ct denotes the cycle threshold. RNA degradation rates were obtained by fitting log-transformed RNA levels within defined time windows. Raw RT-qPCR data are provided in **Supplementary File 1**.

### Data analysis

#### Single-molecule tracking from live and fixed cells

As previously described (20), data was analyzed in Matlab using Oufti (38) for cell detection from bright-field images, u-track (39) for fluorescence spot detection and linking into tracks, a custom-built Matlab code called spotNorm (20) for finding a fluorescence spot’s normalized position in a cell, and custom-built Python code for calculating the membrane-binding percentage (MB%).

##### Oufti

Cell outlines were computed from bright field images using Oufti software developed by Jacobs-Wagner Lab (38). The parameters used for Oufti for mEos3.2 labeled cells are the same as described previously (20).

##### spotNorm

Normalized spot positions within cells were calculated as previously described (20). Briefly, spot locations were normalized along the minor axis (xNorm) and the major axis (yNorm) of each cell. xNorm histograms were constructed using the first four spots per track located within the cylindrical region of the cell. The SEM shown for the xNorm histograms (e.g., **Fig. 2c**-**i**) was estimated by bootstrapping across cells, but it is very small, often smaller than the data line width.

##### u-track for mEos3.2 tracking

Fluorescence spots were detected and compiled into tracks using u-track software developed by Danuser Lab and Jaqaman Lab (39). For mEos3.2 tracking, we used the same parameters as our previous paper (20). A notable parameter in u-track is the max pixel linking distance in the Brownian and directed motion models specifically called the upper bound for the Brownian search radius (in pixels). We set this parameter to 1-3 pixels depending on the preliminary diffusion coefficient based on trajectories identified using u-tracks’ default 5-pixel maximum linking distance, D_5pix_. We previously found that the max linking distance of 5 pixels can create artificial large random jumps in our data set that falsely creates a larger diffusion coefficient. To avoid this error, we set the max linking distance radius as ceiling(2*sqrt(4D_5pix_Δt)), where Δt is the frame time (0.0217 s). Alternatively, we can use 1 pixel if D_5pix_ < 0.0737 µm^2^/s and 2 pixels if 0.0737 µm^2^/s < D_5pix_ < 0.295 µm^2^/s, etc. For proteins with expected slow and fast states (PNPase and enolase), we added one more pixel to prevent cutting out faster diffusing particles. In other words, we used 2 pixels if D_5pix_ < 0.0737 µm^2^/s, 2 pixels if 0.0737 µm^2^/s < D_5pix_ < 0.295 µm^2^/s, etc. Effectively, the max pixel values we used are as follows:

- 5 pixels: Eno (rif)
- 3 pixels: LacZ, LacZ (rif), LacZ (succ), RhlB (rif), Eno, Eno (succ)
- 2 pixels: LacY, LacY (rif), LacY (succ), RNE (succ), RhlB (succ), PNP, PNP (rif), PNP (succ)
- 1 pixel: RNE, RNE (rif), RhlB

##### Diffusion

Tracks lasting at least 12 frames were used for diffusion analysis as previously described (20). Time-averaged (TA) MSD was calculated according to eq. 1, and the diffusion coefficient (D) was calculated by a linear fit using eq. 2 (24):

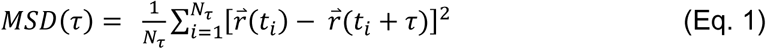

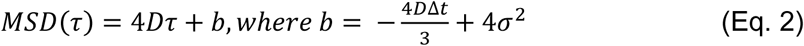

In eq. 1, 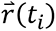 is the 2D vector location of a particle at time *t_i_*, τ is the lag time, and Δt is the number of frames averaged in a track for that lag time. In eq. 2, Δt is the exposure time, and σ is the localization error. The y-intercept, b, is composed of dynamic and static localization errors (24). We fit the first three time points of TA MSD with eq. 2 to get D. This ensures that the fit is done on short time delays (the first 25% of the minimum track length) to avoid MSD at large time intervals with low statistics (40, 41). Once we obtained D from individual tracks, we fit histograms of D.

The Gaussian fitting for a single population (RNE and RhlB) was fit using Eq. 3. For distributions with a long tail (PNP and Eno), a two-population Gaussian was used for fitting (Eq. 4). Fitting was conducted using MATLAB’s function fittype() with reference to either the single-Gaussian equation or double-Gaussian equation saved as a separate function.

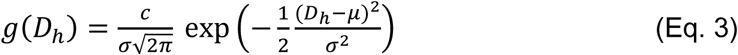

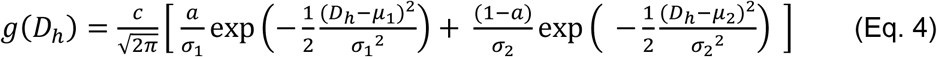

In eq. 3, c is the scaling factor, D_h_ is the diffusion coefficients from D histograms, µ is the mean, and σ is the standard deviation. Similarly, in eq. 4, µ_1_ and µ_2_ are the means of the first and second populations, σ_1_ and σ_2_ are the standard deviations of the first and second populations, a is the fraction of the first population, and (1-a) is the fraction of the second population. As mean values are for diffusion coefficients, we also refer to µ as D_1_ for the single Gaussian fitting and µ_1_ and µ_2_ as D_1_ and D_2_, respectively, for the two-population Gaussian fitting.

### MB% quantification from xNorm histograms

Membrane-bound fraction (MB%) was estimated from xNorm histograms using the modeling framework established in our previous study (20). Briefly, bacterial cells were approximated as cylinders, and molecules were assumed to be homogeneously distributed either on the cell surface (membrane-associated population) or within the cell interior (cytoplasmic population). For each case, the expected xNorm distribution was calculated from the marginal distribution of molecular positions along the short cell axis. The observed distribution was then modeled as a mixture of membrane and cytoplasmic populations, with the membrane-associated fraction corresponding to MB%.

To account for imaging effects, the model incorporated localization uncertainty and finite axial detection range, as described previously (20). Cell width measured from bright-field images was used to scale the model to each dataset, and xNorm values were fit in normalized coordinates. Experimental histograms of absolute xNorm values were fit to the model using Markov chain Monte Carlo (MCMC) sampling with uniform priors on all fitted parameters, including MB%, localization error, dilation factor, and focal cutoff. The posterior estimate of MB% was taken as the membrane-bound fraction. Full mathematical derivation, implementation details, and parameter definitions are provided in (20).

### Data, Materials, and Software Availability

All code used for data analysis is available in Github (sjkimlab/ 2025_RNaseE). Processed microscopy data are available on Zenodo (10.5281/zenodo.20384234) and RT-qPCR data are included in the manuscript as Supplementary File 1.

## Acknowledgements

We thank members of Kim Lab for critical reading of the manuscript. This work was supported by Searle Scholars Program, NIH (R35GM143203), and NSF Science and Technology Center for Quantitative Cell Biology (2243257).

## Author contributions

L.T. and Sangjin Kim designed research; L.T., Y.-H.W., K.W., and Seunghyeon Kim performed research and data analysis; L.T. and Sangjin Kim wrote the manuscript with input from all authors; Sangjin Kim supervised the study and acquired funding.

