## Supplementary information for "Dynamic partitioning shapes the *in vivo* organization of the *E. coli* RNA degradosome"

### Supplementary tables

#### Table S1: Strains used in this study

| Strain number | Genotype | Source | Use |
|---|---|---|---|
| SK1 | MG1655 | | Imaging, Cloning |
| SK98 | MG1655 Δ*lacYA* | CJW5461(1) | RT-qPCR |
| SK187 | MG1655 *rne*:*rne-mEos3.2 kan* | (2) | Imaging, Cloning |
| SK249 | MG1655 *rne*::*rne*ΔMTS-*mEos3.2 kan* | (2) | Imaging |
| SK290 | MG1655 *rne*::*rne-mEos3.2* | (2) | Cloning |
| SK292 | MG1655 *lacYA*::*lacY-mEos3.2 kan* | (2) | Imaging |
| SK301 | MG1655 Δ*lacYA* Δ*rhlB*::*kan* | This study | RT-qPCR |
| SK306 | MG1655 Δ*lacYA* Δ*pnp*::*kan* | This study | RT-qPCR |
| SK345 | MG1655 *pnp*::*pnp-mEos3.2 kan* | This study | Imaging |
| SK350 | MG1655 eno::*eno-mEos3.2 kan* | This study | Imaging |
| SK375 | MG1655 *rhlB*::*rhlB-mEos3.2 kan* | This study | Imaging |
| SK407 | MG1655 *lacZYA::lacZ-mEos3.2 kan* | (2) | Imaging |
| SK421 | MG1655 Δ(*lacZ* 5'UTR)::weakRBS2 Δ*lacYA* | (3) | RT-qPCR |
| SK431 | MG1655 Δ(*lacZ* 5'UTR)::weakRBS2 Δ*lacYA* Δ*pnp*::*kan* | This study | RT-qPCR |
| SK437 | MG1655 Δ(*lacZ* 5'UTR)::weakRBS2 Δ*lacYA* Δ*rhlB*::*kan* | This study | RT-qPCR |

#### Table S2: Strain construction

| Strain number | Construction methods |
|---|---|
| SK301 | The *kan* cassette was amplified from pKD13 using the following primers and integrated into the chromosome of SK98 to replace the *rhlB* gene by lambda Red recombination.<br>rhlB_KO_F:<br>CGGATACGCTTTCGTAAAGCAATAGTAAGCTGATATTCTACCACACTATGATT CCGGGGATCCGTCGACC<br>rhlB_KO_R:<br>TGAATGATTTTGAGTATGACATTTTTTATTTAACCTGAACGACGACGATTTGTA GGCTGGAGCTGCTTCG |
| SK306 | The *kan* cassette was amplified from pKD13 using the following primers and integrated into the chromosome of SK98 to replace the *pnp* gene by lambda Red recombination.<br>pnp_KO_F:<br>CCCGCCGCAGCGGAGGGCAAATGGCAACCTTACTCGCCCTGTTCAGCAGCA TTCCGGGGATCCGTCGACC<br>pnp_KO_R: |

| | |
|---|---|
| | ACACCAGTGCCGTAAGGTACTGTCTAAGAAAGAGAAAGGATATTACATTGTG TAGGCTGGAGCTGCTTCG |
| SK345 | *mEos3.2-kan* region in SK187 was amplified using the primers pnp_mEOS_F and pnp_mEOS_R2 and then integrated into the *pnp* region in MG1655 by lambda Red recombination.<br>pnp_mEOS_F:<br>AGTCTCAACCTGCTGCAGCACCGGAAGCTCCGGCTGCTGAACAGGGCGAG CTCGAGGGTCCGGCTGGTCTGATGTCG<br>pnp_mEOS_R2:<br>ACTCCCGAAGACCACGGTTGAATGAACGTCCTGTTCCCGGTTGCTAACAACT TTGAGCTAATTATCCTTAGTTCC |
| SK350 | *mEos3.2-kan* region in SK187 was amplified using the primers eno_mEOS_F and eno_mEOS_R and then integrated into the *eno* region in MG1655 by lambda Red recombination.<br>eno_mEOS_F :<br>TGGGCGAAAAAGCACCGTACAACGGTCGTAAAGAGATCAAAGGCCAGGCAC TCGAGGGTCCGGCTGGTCTGATGTCG<br>eno_mEOS_R:<br>CCATAAAAAATGCCAGCCCGGAGGCTGGCATTTTTAAATCAGATAAAGTCAG TCCTTTGAGCTAATTATCCTTAGTTCC |
| SK375 | *mEos3.2-kan* region in SK187 was amplified using the primers rhlB_mEOS_F and rhlB_mEOS_R and then integrated into the *rhlB* region in MG1655 by lambda Red recombination.<br>rhlB_mEOS_F:<br>GCAATGGTCCGCGTCGTACTGGCGCTCCGCGTAATCGTCGTCGTTCAGGTC TCGAGGGTCCGGCTGGTCTGATGTCG<br>rhlB_mEOS_R:<br>CTCTTGCCATCTTGATACAGTTTGAATGATTTTGAGTATGACATTTTTTATCTT TGAGCTAATTATCCTTAGTTCC |
| SK431 | Same as SK306, except SK421 strain was used for lambda Red recombination. |
| SK437 | Same as SK301, except SK421 strain was used for lambda Red recombination. |

#### Table S3: Doubling times and cell sizes

| Strain number | Doubling time (min) | Live cell length (μm) | Live cell width (μm) |
|---|---|---|---|
| M9 gly + CAAT | | | |
| SK1 (MG1655) | 82 ± 10 | -- | -- |
| SK187 | 89 ± 9 | 3.36 ± 0.48 | 1.068 ± 0.066 |
| SK345 | 88 ± 3 | 3.26 ± 0.03 | 1.048 ± 0.005 |
| SK350 | 89 ± 2 | 3.26 ± 0.03 | 1.077 ± 0.004 |
| SK375 | 94 ± 5 | 3.28 ± 0.03 | 1.096 ± 0.003 |

| M9 succinate | | | |
|---|---|---|---|
| SK1 (MG1655) | 349 ± 116 | -- | -- |
| SK187 | 223 ± 39 | 3.36 ± 0.02 | 0.970 ± 0.002 |
| SK345 | 266 ± 64 | 3.19 ± 0.03 | 0.957 ± 0.003 |
| SK350 | 302 ± 9 | 3.09 ± 0.04 | 0.957 ± 0.003 |
| SK375 | 325 ± 125 | 3.30 ± 0.02 | 1.003 ± 0.002 |

All numbers are mean ± std. For doubling time, std was calculated from biological replicates. For cell length and width, std was calculated from cell populations.

#### Table S4: RT-qPCR primer sequences

| | **Sequence (5' to 3')** |
|---|---|
| lacZ530F | TTTTACGCGCCGGAGAAAAC |
| lacZ530R | AGTCGGTTTATGCAGCAACG |
| lacZ2732F | TTACTGCCGCCTGTTTTGAC |
| lacZ2732R | TGTAGCGGCTGATGTTGAAC |

#### Table S5: Histogram Gaussian fitting results

| Figure | Protein | # of pop | $R^2$ | c | Population | D ($\mu m^2/s$) | σ ($\mu m^2/s$) |
|---|---|---|---|---|---|---|---|
| 2a | RNE | 1 | 0.98 | 0.010 [0.010, 0.011] | | $D_1$: 0.014 [0.013, 0.016] | $\sigma_1$: 0.014 [0.013, 0.015] |
| 2b | RhlB | 1 | 0.98 | 0.010 [0.009, 0.011] | | $D_1$: 0.014 [0.013, 0.015] | $\sigma_1$: 0.014 [0.012, 0.015] |
| 2c | PNP | 2 | 0.99 | 0.010 [0.0098, 0.0103] | $P_1$: 0.56 [0.47, 0.65]<br>$P_2$: 0.44 [0.35, 0.53] | $D_1$: 0.014 [0.014, 0.015]<br>$D_2$: 0.038 [0.032, 0.044] | $\sigma_1$: 0.014 [0.012, 0.014]<br>$\sigma_2$: 0.029 [0.026, 0.032] |
| 2d | Eno | 2 | 0.99 | 0.0090 [0.0088, 0.0093] | $P_1$: 0.60 [0.56, 0.65]<br>$P_2$: 0.40 [0.35, 0.44] | $D_1$: 0.014 [0.014, 0.015]<br>$D_2$: 0.043 [0.038, 0.049] | $\sigma_1$: 0.014 [0.014, 0.015]<br>$\sigma_2$: 0.041 [0.038, 0.044] |
| S2a | RNE, M9 succinate | 1 | 0.97 | 0.0099 [0.0094, 0.0104] | | $D_1$: 0.025 [0.024, 0.026] | $\sigma_1$: 0.023 [0.022, 0.024] |

| | | | | | | | |
|---|---|---|---|---|---|---|---|
| S2b | RhlB,<br>M9 succinate | 1 | 0.96 | 0.0096<br>[0.0090,<br>0.0102] | | $D_1$: 0.022<br>[0.021, 0.024] | $\sigma_1$: 0.023<br>[0.021,<br>0.024] |
| S2c | PNPase,<br>M9 succinate | 2 | 0.99 | 0.0101<br>[0.0099,<br>0.0102] | $P_1$: 0.54<br>[0.48, 0.59]<br><br>$P_2$: 0.46<br>[0.41, 0.52] | $D_1$: 0.021<br>[0.021, 0.022]<br><br>$D_2$: 0.056<br>[0.051, 0.061] | $\sigma_1$: 0.018<br>[0.017,<br>0.018]<br>$\sigma_2$: 0.039<br>[0.037,<br>0.041] |
| S2d | Enolase,<br>M9 succinate | 2 | 0.98 | 0.0094<br>[0.0092,<br>0.0097] | $P_1$: 0.58<br>[0.55, 0.62]<br><br>$P_2$: 0.42<br>[0.38, 0.45] | $D_1$: 0.023<br>[0.022, 0.023]<br><br>$D_2$: 0.084<br>[0.075, 0.093] | $\sigma_1$: 0.020<br>[0.020,<br>0.021]<br>$\sigma_2$: 0.072<br>[0.067,<br>0.078] |

See Eq. 3-4 for fitting parameters. Brackets represent the 95% confidence interval (CI).

#### Table S6: Figure data statistics

| Figure number | Strain (SK#) | Strain (description) | Number of tracks or spots | Number of cells |
|---|---|---|---|---|
| 2c | SK292 | LacY | 199,228 spots | 161 |
| 2d | SK407 | LacZ | 218,120 spots | 247 |
| 2e | SK187 | RNE | 143,700 spots | 179 |
| 2f | SK375 | RhlB | 121,420 spots | 200 |
| 2g | SK345 | PNP | 186,108 spots | 221 |
| 2h | SK350 | Eno | 141,748 spots | 211 |
| 3a | SK187 | RNE | 11,260 tracks | 178 |
| 3b | SK375 | RhlB | 9,540 tracks | 198 |
| 3c | SK345 | PNP | 15,098 tracks | 221 |
| 3d | SK350 | Eno | 6,344 tracks | 199 |
| 4a | SK292 | LacY | 199,228 spots | 161 |
| | SK292 | LacY + rif | 73,784 spots | 166 |
| 4b | SK407 | LacZ | 218,120 spots | 247 |
| | SK407 | LacZ + rif | 163,560 spots | 370 |
| 4c | SK187 | RNE | 143,700 spots | 179 |
| | SK187 | RNE + rif | 174,000 spots | 291 |
| 4d | SK375 | RhlB | 121,420 spots | 200 |
| | SK375 | RhlB + rif | 134,088 spots | 273 |
| 4e | SK345 | PNP | 186,108 spots | 221 |
| | SK345 | PNP + rif | 43,988 spots | 257 |
| 4f | SK350 | Eno | 141,748 spots | 211 |
| | SK350 | Eno + rif | 1,026,976 spots | 449 |

| | | | | |
|---|---|---|---|---|
| 5a | SK292 | LacY in M9 gly CAAT | 199,228 spots | 161 |
| | SK292 | LacY in M9 succinate | 500,972 spots | 589 |
| 5b | SK407 | LacZ in M9 gly CAAT | 218,120 spots | 247 |
| | SK407 | LacZ in M9 succinate | 108,388 spots | 243 |
| 5c | SK187 | RNE in M9 gly CAAT | 143,700 spots | 179 |
| | SK187 | RNE in M9 succinate | 388,632 spots | 693 |
| 5d | SK375 | RhlB in M9 gly CAAT | 121,420 spots | 200 |
| | SK375 | RhlB in M9 succinate | 324,412 spots | 590 |
| 5e | SK345 | PNP in M9 gly CAAT | 186,108 spots | 221 |
| | SK345 | PNP in M9 succinate | 189,844 spots | 327 |
| 5f | SK350 | Eno in M9 gly CAAT | 141,748 spots | 211 |
| | SK350 | Eno in M9 succinate | 276,948 spots | 351 |
| S2a | SK187 | RNE (all) | 143,700 spots | 179 |
| | SK187 | RNE (12+) | 108,540 spots | 179 |
| S2b | SK375 | RhlB (all) | 121,420 spots | 200 |
| | SK375 | RhlB (12+) | 92,952 spots | 200 |
| S2c | SK345 | PNP (all) | 186,108 spots | 221 |
| | SK345 | PNP (12+) | 148,248 spots | 221 |
| S2d | SK350 | Eno (all) | 141,748 spots | 211 |
| | SK350 | Eno (12+) | 58,560 spots | 211 |
| S4 | SK407 | LacZ | 270,164 spots | 248 |
| | SK292 | LacY | 262,788 spots | 161 |
| | SK187 | RNE | 177,584 spots | 179 |
| | SK375 | RhlB | 156,732 spots | 201 |
| | SK345 | PNP | 224,140 spots | 223 |
| | SK350 | Eno | 177,960 spots | 211 |
| | SK407 | LacZ + rif | 197,176 spots | 370 |
| | SK292 | LacY + rif | 97,592 spots | 166 |
| | SK187 | RNE + rif | 213,440 spots | 291 |
| | SK375 | RhlB + rif | 190,448 spots | 413 |
| | SK345 | PNP + rif | 52,268 spots | 257 |
| | SK350 | Eno + rif | 1,268,004 spots | 486 |
| | SK407 | LacZ in M9 succinate | 131,324 spots | 243 |
| | SK292 | LacY in M9 succinate | 632,472 spots | 589 |
| | SK187 | RNE in M9 succinate | 395,444 spots | 693 |
| | SK375 | RhlB in M9 succinate | 393,268 spots | 590 |
| | SK345 | PNP in M9 succinate | 228,788 spots | 327 |
| | SK350 | Eno in M9 succinate | 313,100 spots | 351 |
| S5a | SK187, M9 succinate | RNE, M9 succinate | 32,058 tracks | 660 |
| S5b | SK375, | RhlB, M9 succinate | 28,583 tracks | 556 |

| | | | | |
|---|---|---|---|---|
| | M9 succinate | | | |
| S5c | SK345,<br>M9 succinate | PNP, M9 succinate | 17,219 tracks | 308 |
| S5d | SK350,<br>M9 succinate | Eno, M9 succinate | 15,949 tracks | 337 |

#### Supplementary figures

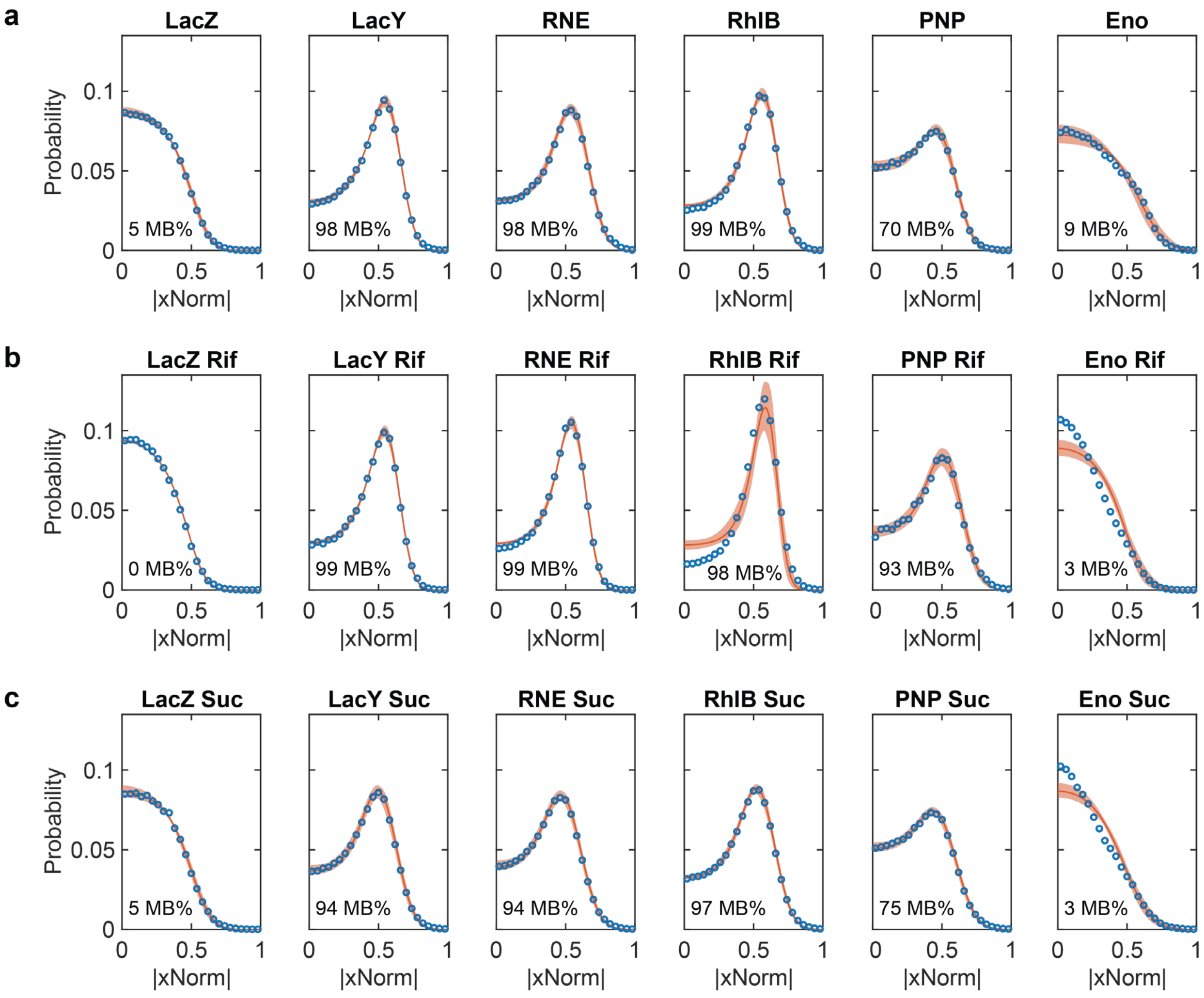

**Figure S1. MB% modeling of xNorm histograms.**
xNorm histograms of control proteins (LacY and LacZ) and RNAD proteins under the reference condition (**a**), after rifampicin treatment (**b**) and in M9 succinate medium (**c**) were analyzed for MB%. In all panels, experimental data (blue circles) is compared with MCMC-based fit (red line). The red shaded region represents the expected range of xNorm histograms generated from parameter values within one standard deviation of the best-fit estimate. The MB% shown in the lower left corner of each plot corresponds to the posterior mean estimated from the MCMC samples.

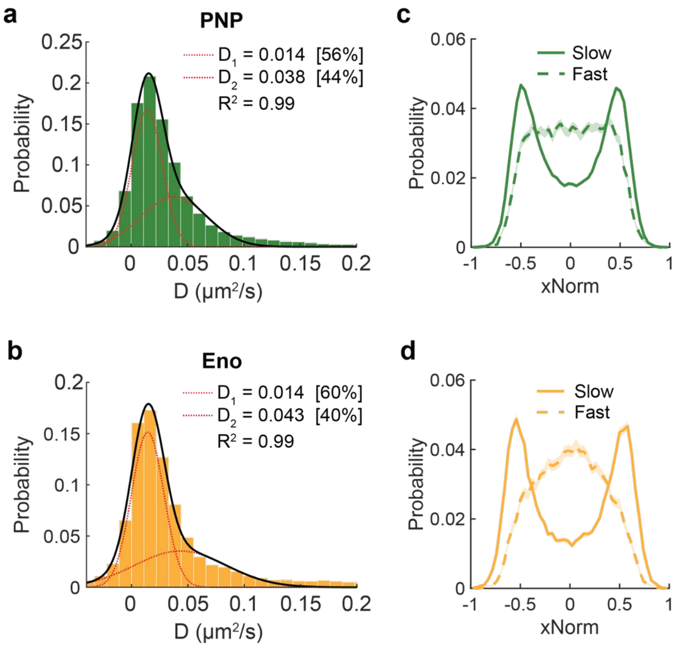

**Figure S2. xNorm histograms of fast and slow PNP and Eno populations.**

(**a**, **b**) Distributions of diffusion coefficients of PNP (a) and Eno (b) with a two-population fit, as shown in Fig. 3c, d. (**c**, **d**) xNorm histograms of slow and fast trajectories of PNP (c) and Eno (d). Slow trajectories were defined as those with D < 0.06 μm$^2$/s, whereas fast trajectories were defined as those with D ≥ 0.06 μm$^2$/s. This diffusion coefficient cutoff was chosen based on the D cut off was chosen based on the upper bound of the fitted slow-population, $D_1$.

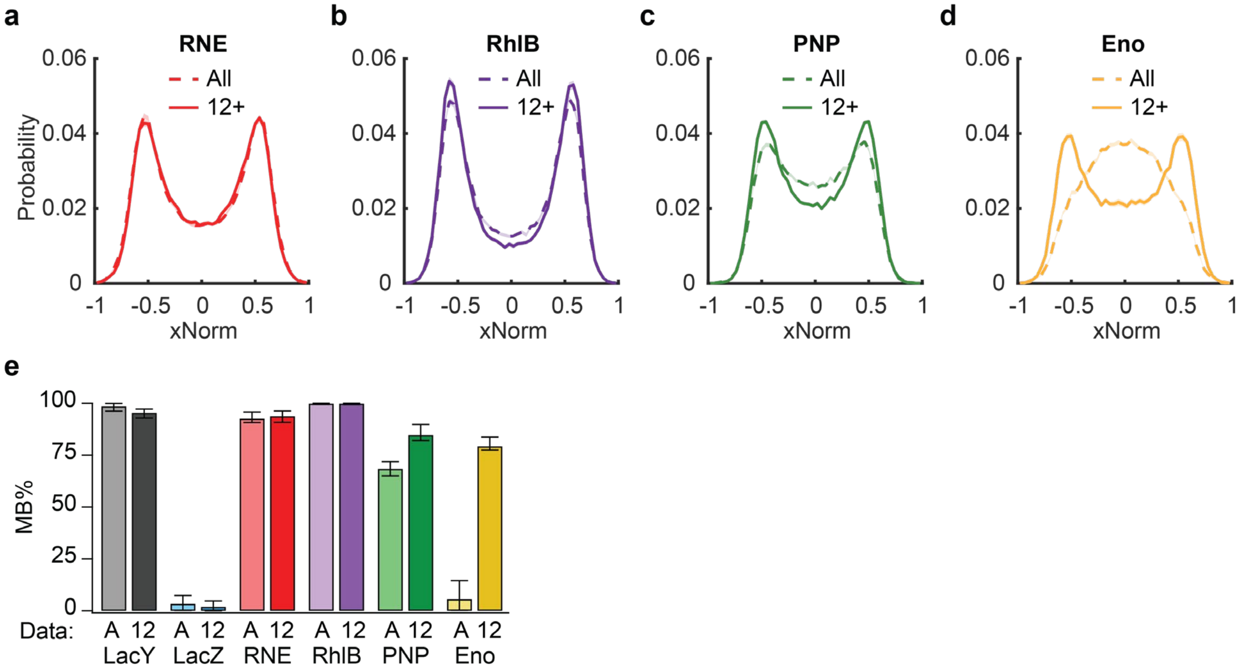

**Figure S3. xNorm histogram analysis of long trajectories used for diffusion measurements.** (**a-d**) xNorm histograms for proteins imaged in live cells, including RNE (a), RhlB (b), PNP (c), and Eno (d). 'All' indicates xNorm distributions calculated from all trajectories, whereas '12+' indicates those calculated only from trajectories at least 12 frames long. (**e**) MB% estimated from the xNorm histograms. 'A' denotes all trajectories, and '12' stands for trajectories with at least 12 frames. Error bars represent 95% CI.

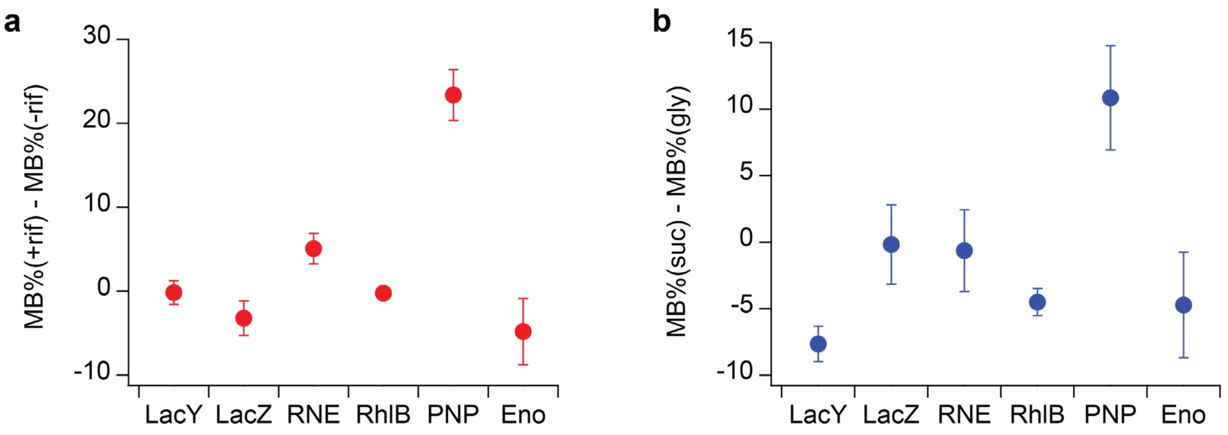

**Figure S4. Posterior estimates of MB% changes in RNAD components across conditions.** (**a**) Changes in MB% after rifampicin treatment, shown as MB%(+rif) - MB%(−rif). (**b**) Differences in MB% between growth conditions, M9 succinate and M9 gly + CAAT, shown as MB%(suc) – MB%(gly), for reference proteins (LacY and LacZ) and RNAD components (RNE, RhlB, PNP, and Eno). Symbols represent the mean of posterior difference distributions generated from MCMC samples, and error bars indicate standard deviations.

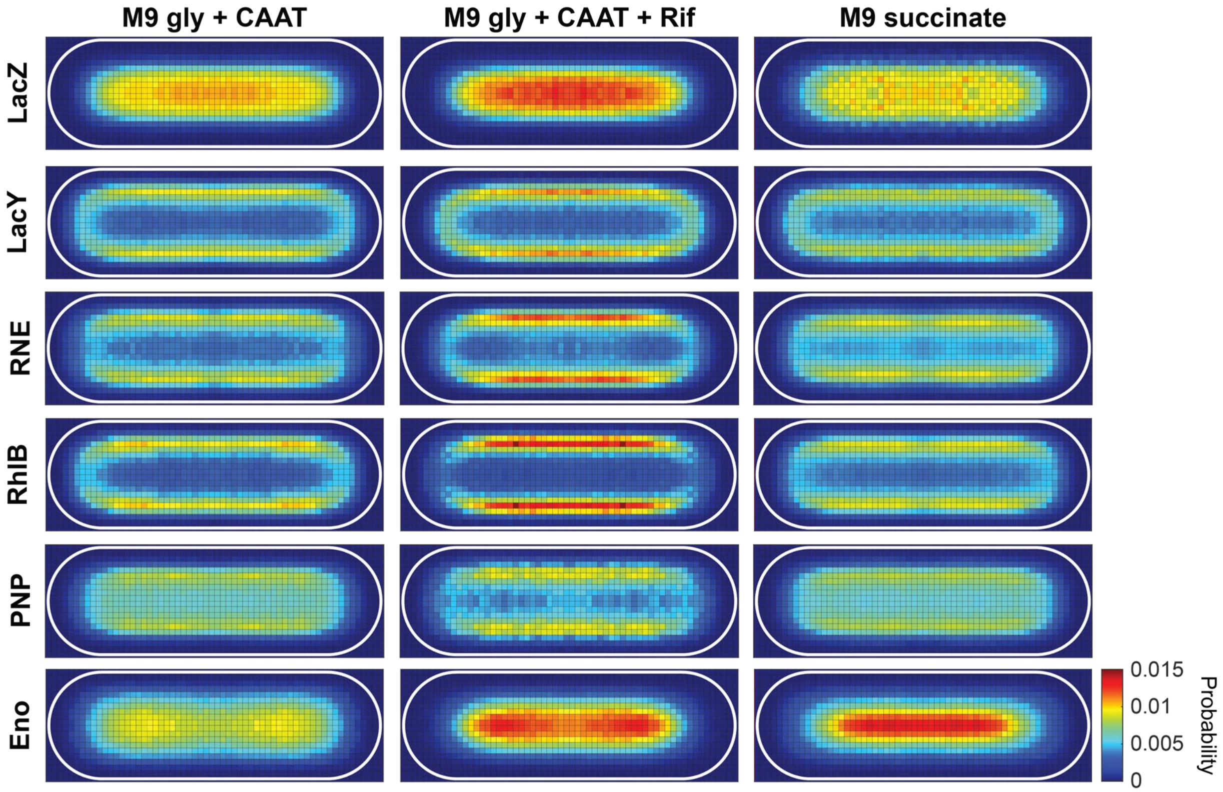

**Figure S5. Two-dimensional histograms of protein localization.**
Color indicates the probability of the molecule within each spatial bin. Subcellular localizations were normalized in the first quartile, expressed as xNorm and yNorm coordinates, and mirrored across the cell axes. Bin size: 48-56 nm. White outlines indicate cell boundaries. The same color scale was used for all histograms. Data statistics are provided in **Table S6**.

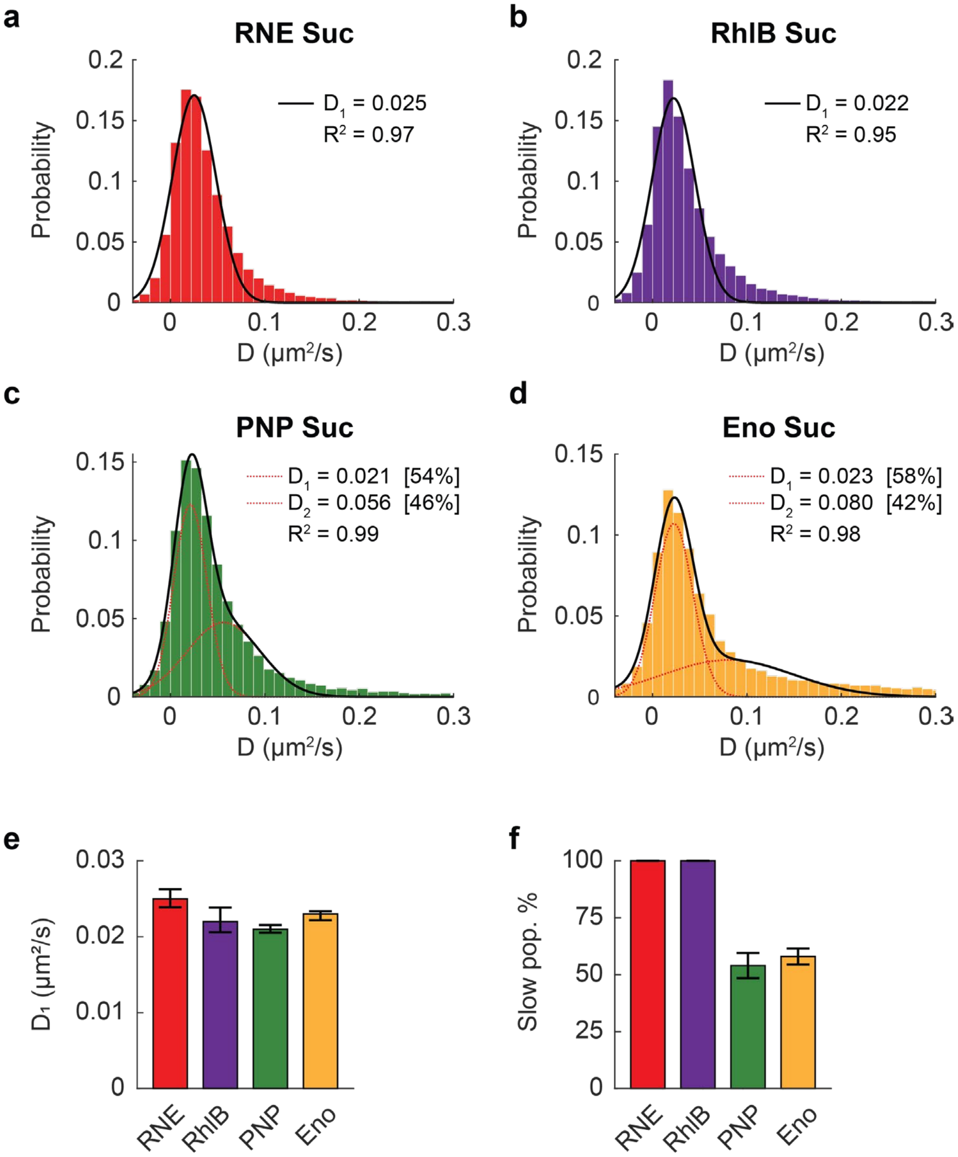

**Figure S6. Diffusion of RNAD component proteins in cells grown in M9 succinate.**
(**a-d**) Distributions of diffusion coefficients for RNE (a), RhlB (b), PNP (c), and Eno (d). The histograms were fit with either a one-Gaussian population model (a,b) or a two-Gaussian population model (c,d). Additional fitting details are provided in **Table S5**. (**e**) $D_1$ values for the RNAD component proteins obtained from the Gaussian fits in (a-d). (**f**) Percentage of the slow population from histogram fitting, with single-population fits treated as 100% slow population. Error bars in (e) and (f) indicate the 95% CI from fitting. Data statistics are provided in **Table S6**.
